# Optimizing transcriptome-based synthetic lethality predictions to improve targeted therapy / immunotherapy treatment prioritization in non-metastatic breast cancer: BC-SELECT

**DOI:** 10.1101/2024.08.15.608073

**Authors:** Yewon Kim, Matthew Nagy, Rebecca Pollard, Padma Sheila Rajagopal

**Affiliations:** Cancer and Data Science Laboratory, Center for Cancer Research, National Cancer Institute, Bethesda, MD, USA; Boston Children’s Hospital, Boston, MA, USA; Women’s Malignancies Branch, Center for Cancer Research, National Cancer Institute, Bethesda, MD, USA

**Keywords:** Synthetic lethality, treatment prediction, breast cancer, immunotherapy, PARP inhibitors

## Abstract

**Background:** Patients with non-metastatic triple-negative (TNBC) or HER2-positive (HER2+) breast cancer do not have tools to predict treatment response or prioritize options prior to starting therapy. Pathologic complete response (pCR) is a proxy that requires neoadjuvant treatment. BC-SELECT leverages genetic interactions (e.g. synthetic lethality (SL)/rescue (SR)) to predict treatment response via pCR from breast cancer gene expression data. BC-SELECT involves (1) “gene-pair identification” that identifies clinically relevant partner genes from large-scale datasets for a given targeted therapy or immunotherapy target, and (2) “training / parameter tuning” using clinical trials to predict treatment response. We evaluated BC-SELECT’s ability to predict pCR for trastuzumab, poly (ADP-ribose) polymerase (PARP) inhibitors, and immunotherapy, with tuning supported by three trials (n=86) and validation on nine unseen trials (n=722).

**Results:** BC-SELECT significantly predicted pCR in 6/9 trials, including all PARP inhibitors (ROC-AUCs 0.6-0.8; Odds Ratio (OR): 1.9-3.5) and immunotherapy (ROC-AUCs 0.7-0.8; OR: 3.2-7.1). BC-SELECT significantly distinguished between pCR and residual disease in 8/9 trials.

**Conclusions:** No current standard-of-care, pre-treatment approaches predict benefit in non-metastatic TNBC and HER2+ breast cancer. BC-SELECT leverages genetic interactions to predict treatment response from gene expression data. Our findings support development of BC-SELECT towards prioritizing PARP inhibitors and immunotherapy in non-metastatic breast cancer.

## BACKGROUND

Gene expression data has transformed clinical management in non-metastatic breast cancer. Patients with early-stage hormone-receptor positive (HR+) disease undergo testing to predict chemotherapy benefit^1–9^. There is no equivalent test for targeted therapy used in HR+ breast cancer (such as CDK4/6 inhibitors). HER2+ or TNBC also lack predictive gene expression tests and require neoadjuvant therapy and surgery to assess pCR to guide adjuvant treatment.

Patients with HER2+ or TNBC may have competing treatment options across neoadjuvant/adjuvant settings, creating a need for predictors. For example, adjuvant PARP inhibitors are approved to treat patients with “high-risk early breast cancer” due to inherited *BRCA1/2* pathogenic variants (most commonly TNBC), and neoadjuvant/adjuvant immunotherapy is approved for “high-risk, early-stage TNBC.”^10,11^ Multiple clinical trials in metastatic breast cancer (KEYLYNK-009, MEDIOLA, PEMBRACA) fail to show an outcome benefit for combining these therapies. The lack of data in nonmetastatic breast cancer is described explicitly in current NCCN guidelines.^12–16^ When multiple targeted therapies / immunotherapies are possible, sequential ordering may offer the same benefits without the toxicity of combinations.

Many published computational approaches demonstrate the prognostic and potentially predictive potential of gene expression in non-metastatic TNBC or HER2+ breast cancer, but these approaches must still navigate current clinical challenges^17^. Recent studies support “immune-related” gene signatures to predict immunotherapy response^18–20^. HER2 gene expression is evolving alongside immunohistochemistry and FISH^21,22^. Broader transcriptomic data can now be routinely obtained on commercial testing and identifies not only fusions for FDA-approved therapeutic indications, but also gene expression levels, positioning it as a real potential biomarker.^23,24^

Synthetic lethality (SL), a form of genetic interaction in which two genes must be concurrently inactivated to cause cell death, is a mechanism of action with known clinical relevance in breast cancer^25,26^. PARP inhibitors are approved to treat breast cancers with inherited *BRCA1/2* pathogenic mutations due to the synthetic lethality between *BRCA1/2* and *PARP*1/2/3^10,27,28^. Other genetic interactions, such as synthetic dosage lethality (SDL), in which one gene must be over-expressed and another under-expressed^29^, or synthetic rescue (SR), in which inactivation of one gene that would cause cell death is ameliorated by altered activation of another gene, are being actively studied for translational and clinical application^30^.

Computational approaches to identify genetic interactions through gene expression have been recently developed. SELECT^31^ uses large-scale cell-line and pan-cancer patient tumor transcriptomic data to identify clinically relevant SL/SR partners for cancer therapies. A commercial adaptation, ENLIGHT^32^, focuses on advanced cancers. Both algorithms predict treatment response in retrospective clinical trials and demonstrate SR’s ability to predict immunotherapy response^31,32^. However, their testing in non-metastatic breast cancer was limited. Evaluated breast cancer therapies included only endocrine therapy or trastuzumab without other targeted therapies or immunotherapy.

Accordingly, we present BC-SELECT, designed to facilitate prediction of response to current and future targeted therapy and immunotherapy in non-metastatic breast cancer. We evaluate BC-SELECT’s ability to predict pCR for trastuzumab, PARP inhibitors, and immunotherapy by training/tuning on three trials (86 patients) and validating on nine unseen trials (722 patients). BC-SELECT predicts treatment response for PARP inhibitors – not previously shown by SELECT or ENLIGHT – and immunotherapy across breast cancer subtypes, positioning it as a tool to eventually prioritize targeted therapy and immunotherapy in non-metastatic breast cancer.

## RESULTS

### Overview of the BC-SELECT framework and adaptations to predict pCR for therapeutic options in non-metastatic breast cancer

Figure 1 shows the BC-SELECT pipeline. BC-SELECT has two parts: “Gene-pair identification” and “training / parameter tuning.” We used large-scale cell-line and breast cancer datasets to find clinically relevant, interacting gene pairs A and B (in which gene A is the target(s) for targeted therapy or immunotherapy, and multiple partner B genes are identified). Expression levels of partner B genes were used to generate a patient’s predicted response score to a treatment targeting gene A. We trained and tuned parameters using one clinical trial that reported pCR per therapy (Supplementary Table 1, Supplementary Methods). We validated BC-SELECT using nine unseen, independent clinical trials in patients with stage I-III breast cancer: Four using trastuzumab in HER2+ disease, three using PARP inhibitors in TNBC, and two using PD-1/PD-L1 immunotherapy in TNBC (Supplementary Table 1). Three trials also recruited HR+ patients. Patients’ responses were evaluated only within their own clinical trials to avoid batch effect or inaccurate cross-dataset normalization.

**Figure 1:**
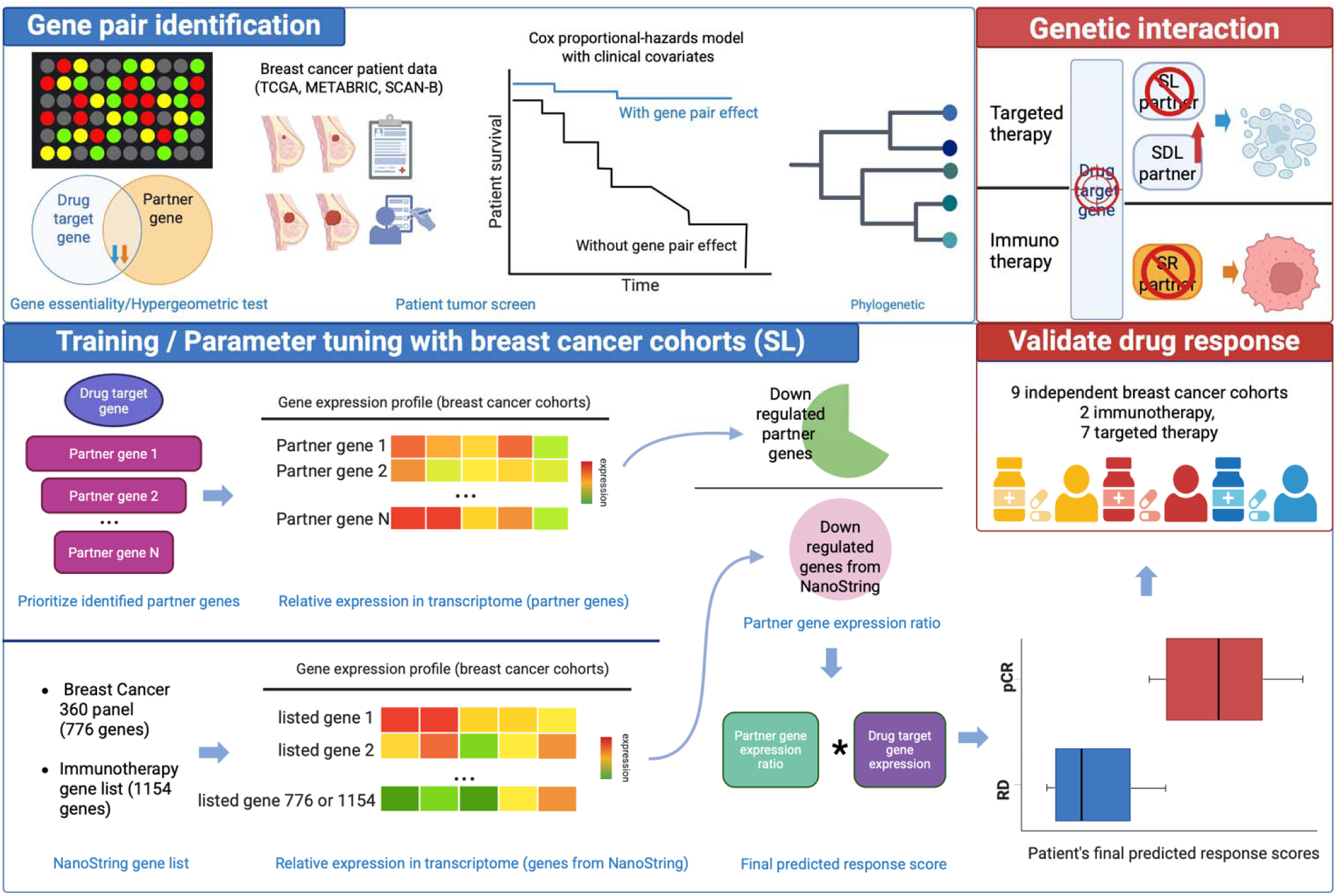
A graphical overview illustrating the adaptation of BC-SELECT pipeline. BC-SELECT involves two steps (in blue): gene-pair identification and training / parameter tuning. The first step focuses on identifying gene pairs involved in genetic interactions. We then adjust parameters to optimize drug response prediction performance using a subset of available clinical trials during the second step. We are then able to apply this to unseen clinical trials and did so in nine additional validation breast cancer clinical trials.

### Partner gene biology

Partner genes identified by BC-SELECT for therapeutic targets (HER2, PARP1/2, and PD-1/PD-L1) (Supplementary Table 2) are associated with complementary pathways. Significant partner pathways (p<0.01) for HER2 included microtubule binding (n=10 genes) and tubulin binding (n=10). Significant partner pathways for PARP included ATP hydrolysis activity (n=7) and helicase activity (n=5). Significant partner pathways (p<0.01) for PD-1/PD-L1 were related to MHC protein complex binding (n=2-4). *FLT3, PLOD1, TRIB2, and TK1* overlapped between SDL/trastuzumab partners and SL/SDL/PARP inhibitor partners. No partner genes were shared between targeted therapy and immunotherapy.

### BC-SELECT significantly predicted pCR in response to PARP inhibitors and immunotherapy

Across all trials, complete responders (pCR = 1) demonstrated consistently higher predicted scores from BC-SELECT compared to those with residual disease/non-pCR (pCR = 0).

We derived odds ratios between pCR and non-pCR based on BC-SELECT final predicted response scores (Figure 2). Odds ratios were statistically significant in 6/9 trials using a threshold of 0.51, including all PARP inhibitor and all immunotherapy trials (Table 1). We observed a statistically significant difference between final predicted response scores with pCR compared to non-pCR for both targeted therapy (6/7 trials) (Figure 3A) and immunotherapy (2/2 trials) (Figure 3B) (FDR-corrected Wilcoxon rank-sum test).

**Figure 2:**
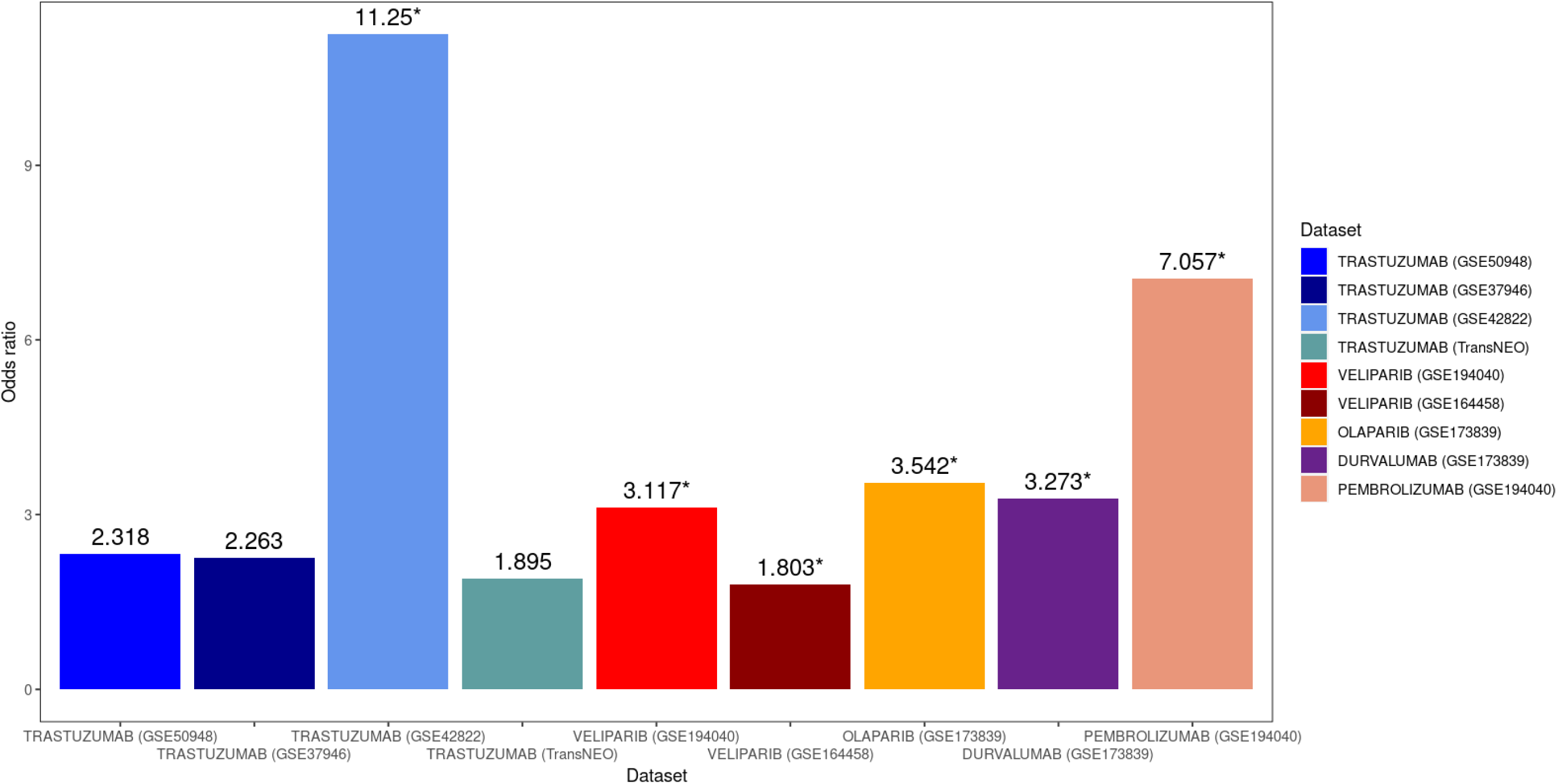
Odds ratios between pCR and non-pCR outcome using final predicted response scores from BC-SELECT. X-axis shows the specific clinical trial, and Y-axis shows the odds ratio. Using a threshold of 0.51, we calculated the odds ratios between pCR and non-pCR outcome across 9 breast cancer clinical trials. Trastuzumab is shown in blue/green, PARP inhibitors in red/orange, and immunotherapy in purple/pink. All odds ratios were greater than 1, and prediction in 6 trials showed statistical significance (marked with *), with the 95% confidence interval not containing 1. Detailed 95% confidence intervals of odds ratios and multiplicity-corrected p-values from Fisher exact tests are provided in Table 1.

**Figure 3:**
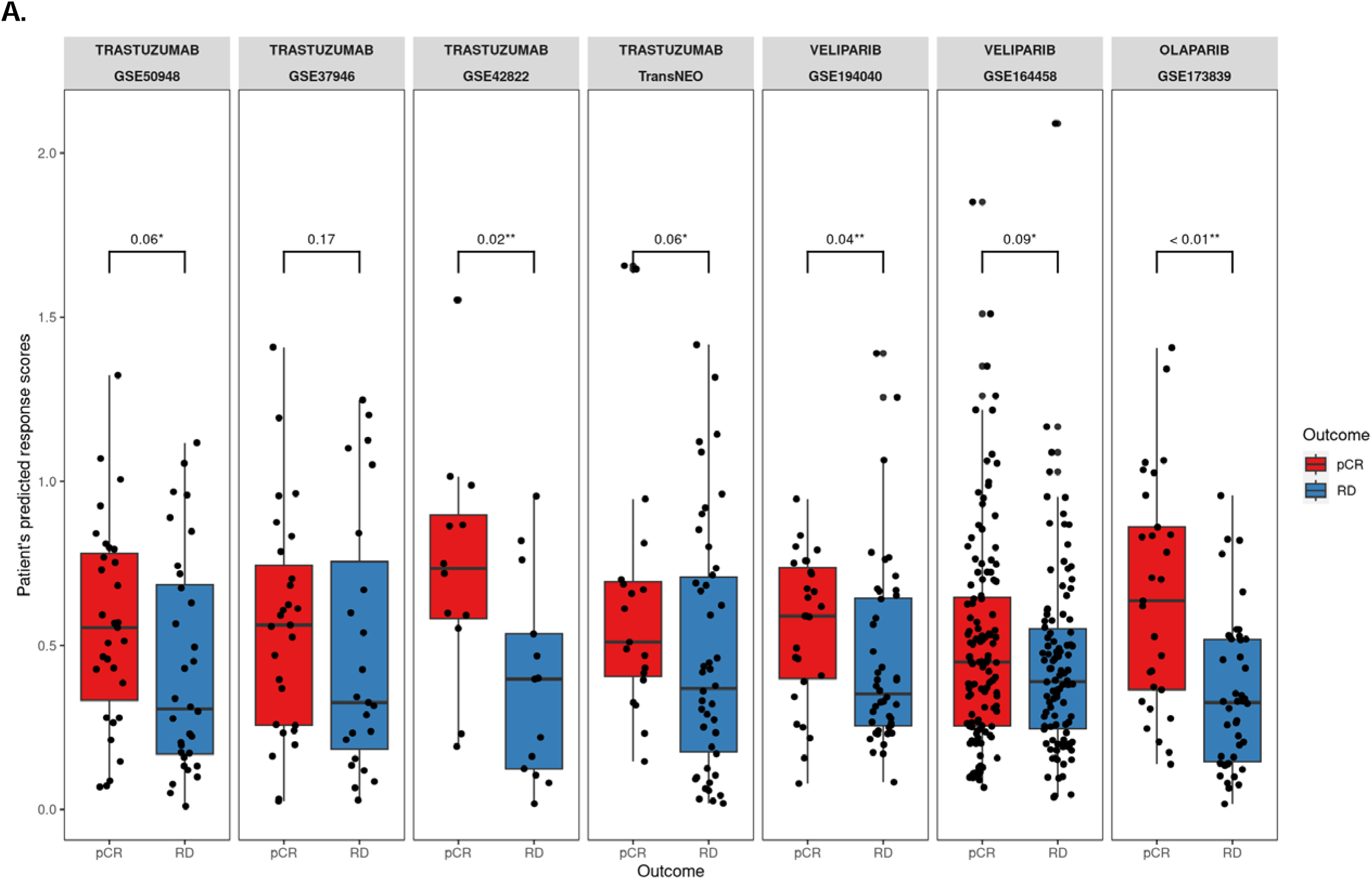
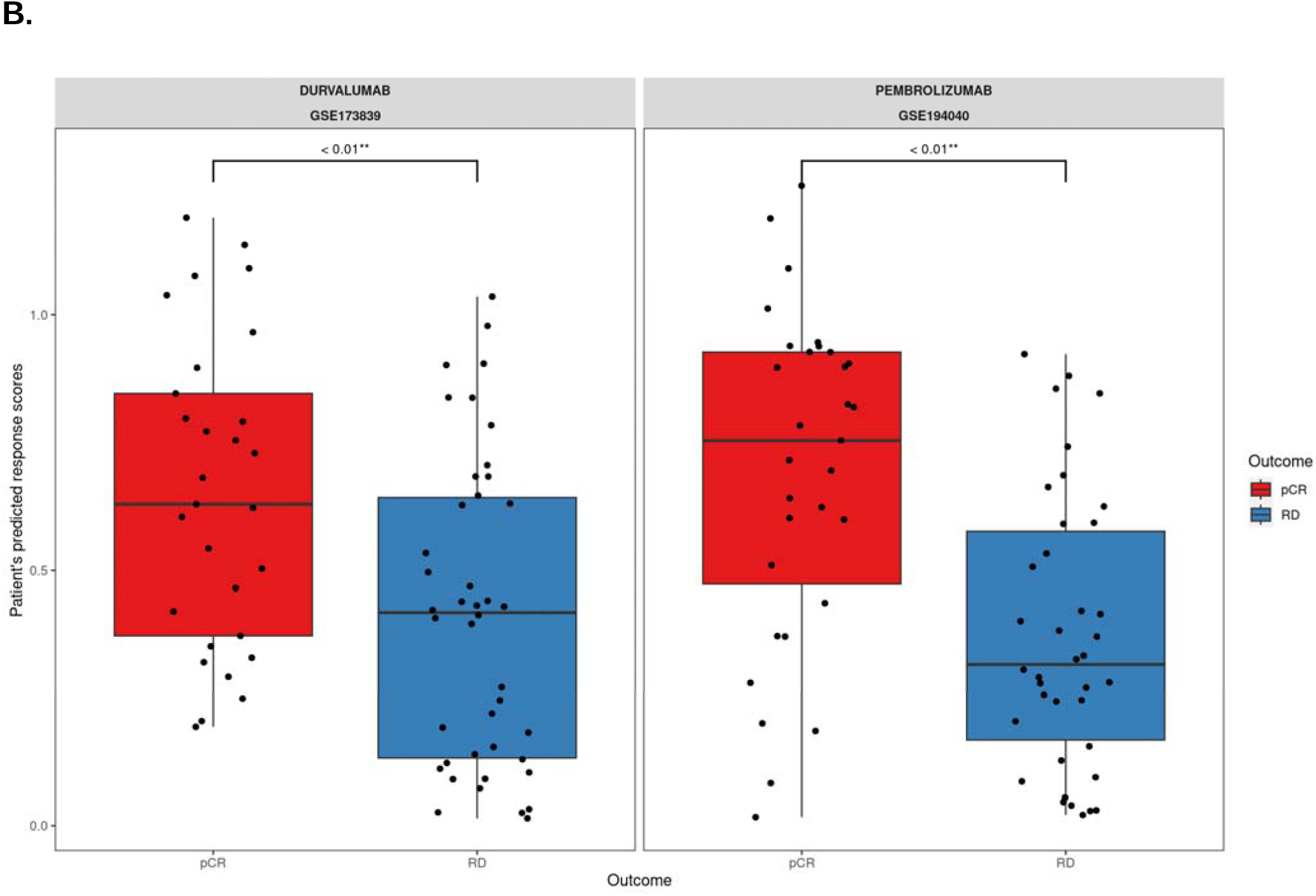
Patient’s final predicted response scores of BC-SELECT among pCR versus non-pCR outcome. Red = pCR, Blue = RD: Residual disease/non-pCR outcome. **A. Targeted therapies.** X-axis shows the specific clinical trial, and Y-axis shows the distribution of predicted response scores within the trial. Final predicted BC-SELECT scores are significantly higher in pCR compared with RD in 6 out of 7 trials. **B. Immunotherapy.** Final predicted BC-SELECT scores are also significantly higher in pCR versus RD in both immunotherapy trials. Significance was determined via the Wilcoxon rank-sum test with correction for multiple hypothesis testing. Adjusted p-values are displayed in each box plot, where p < 0.1 is indicated by a single asterisk (*) and p < 0.05 by a double asterisk (**).

**Table 1:** Odds ratios between pCR and non-pCR outcome using patients’ final predicted response scores from BC-SELECT. pCR: Pathologic complete response. RD: Residual disease/non-pCR outcome. SL: Synthetic Lethality. SDL: Synthetic Dosage Lethality. SR: Synthetic Rescue. All odds ratios were greater than 1, and six trials showed statistical significance (bold) based on 95% confidence interval. Adjusted p-values as determined through the Fisher’s exact test and multiple hypothesis correction are displayed in the last column, where p < 0.1 is indicated by a single asterisk (*) and p ≤ 0.05 by a double asterisk (**).

|  | DRUG | TARGET | TRIAL | GENETIC INTERACTION | Odds ratio | 95% Confidence Interval | Fisher exact test adjusted P-value |
| --- | --- | --- | --- | --- | --- | --- | --- |
| Immuno therapy | DURVALUMAB | PDCD1, CD274 | GSE173839<sup>66</sup><br>(pCR:29, RD:42) | SR | **3.273** | **(1.22, 8.782)** | **0.03\*\*** |
|  | PEMBROLIZUMAB |  | GSE194040<sup>20</sup><br>(pCR:31, RD:38) | SR | **7.057** | **(2.428, 20.514)** | **< 0.01\*\*** |
| Targeted therapy | VELIPARIB | PARP1, PARP2 | GSE194040<sup>20</sup><br>(pCR: 27, RD: 44) | SDL | **3.117** | **(1.151, 8.438)** | **0.07\*** |
|  |  |  | GSE164458<sup>33</sup><br>(pCR:127, RD:110) | SL | **1.803** | **(1.049, 3.098)** | **0.07\*** |
|  | OLAPARIB | PARP1, PARP2, PARP3 | GSE173839<sup>66</sup><br>(pCR:29, RD:42) | SL | **3.542** | **(1.307, 9.6)** | **0.05\*\*** |
|  | TRASTUZUMAB | ERBB2 (HER2) | GSE50948<sup>62</sup><br>(pCR:31, RD:32) | SDL | 2.318 | (0.839, 6.404) | 0.18 |
|  |  |  | GSE37946<sup>63</sup><br>(pCR:27, RD:23) | SDL | 2.263 | (0.727, 7.047) | 0.28 |
|  |  |  | GSE42822<sup>64</sup><br>(pCR:12, RD:13) | SDL | **11.25** | **(1.647, 76.852)** | **0.05\*\*** |
|  |  |  | TransNEO<sup>65</sup><br>(pCR:19, RD:46) | SL | 1.895 | (0.643, 5.589) | 0.28 |

BC SELECT’s final predicted response score outperformed its two components: target gene expression (gene A) and partner gene expression ratio (genes B) (Table 2). For targeted therapy, BC-SELECT final predicted response scores predicted pCR significantly better than chance (one-tailed empirical p-value < 0.025) in four trials (2/3 trials with a PARP inhibitor and 2/4 trastuzumab trials), with AUCs ranging from 0.63-0.8, while partner gene expression ratio and target gene expression were each statistically significant in two trials. For immunotherapy, BC-SELECT and partner gene expression ratio were both statistically significant for both trials in predicting response compared to chance, but ROC-AUCs were higher with BC-SELECT (0.71,0.78 vs. 0.62,0.66). Target gene expression was significant in one of two trials. Precision-recall curves are shown in Supplementary Figure 1.

**Table 2:**
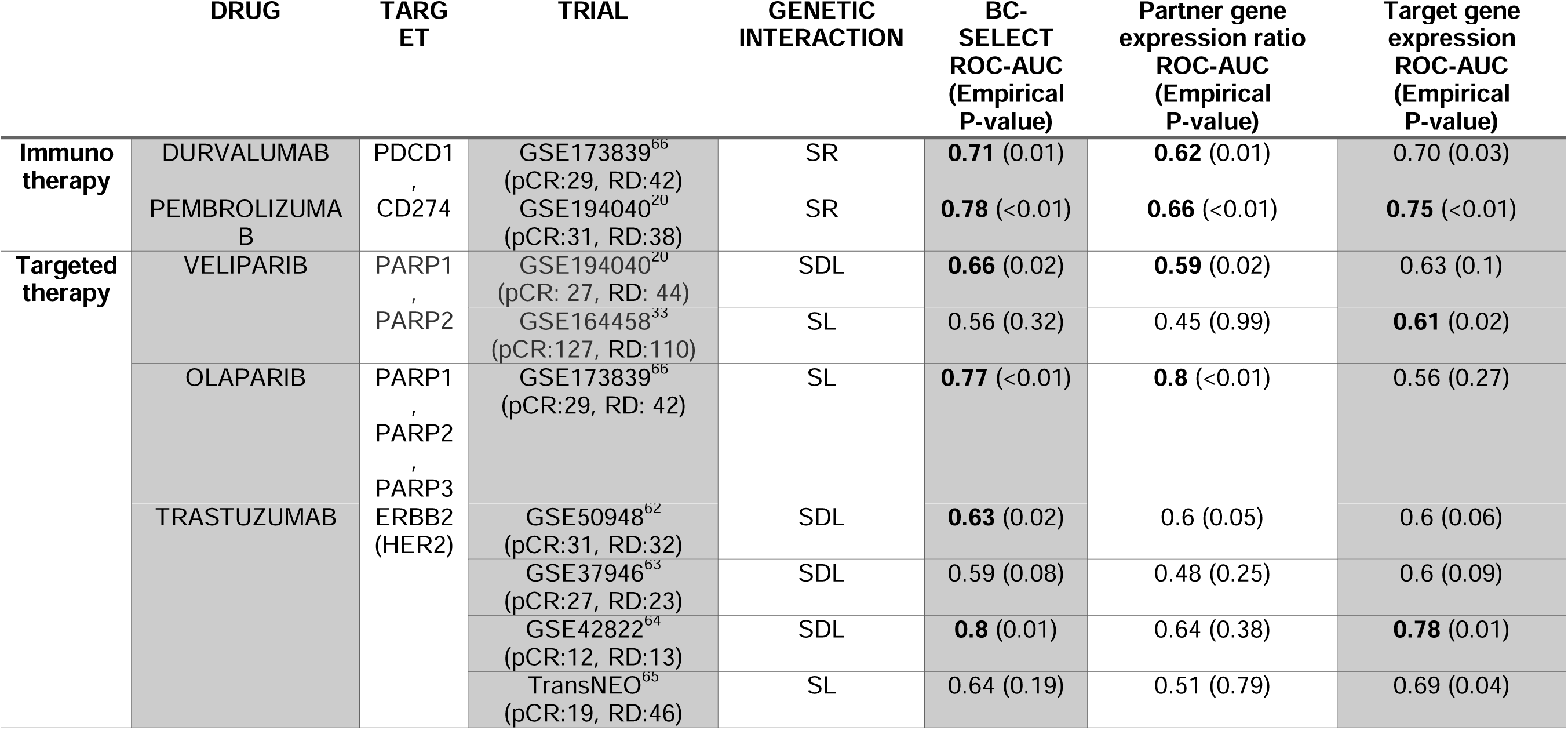
ROC-AUC values and empirical p-values for BC-SELECT and its score components (partner gene expression ratio and target gene expression) in predicting nine breast cancer clinical trials. pCR: Complete responders. RD: Residual disease/non-pCR outcome. SL: Synthetic Lethality. SDL: Synthetic Dosage Lethality. SR: Synthetic Rescue. We report ROC-AUCs and empirical p-values, which describe the performance within a Monte Carlo-generated null distribution. ROC-AUC values are highlighted in bold if the ROC-AUC values fall within the top 2.5% of null AUC distributions. For targeted therapies, the reported ROC-AUC value is the higher one that was generated using either SL or SDL.

To confirm that BC-SELECT scores were not inadvertently increased as a factor of being assigned/given targeted therapy, we compared the theoretical AUC of PARP inhibitor response for both placebo arms in BrighTNess (GSE164458)^33^ (B and C, 245 patients, SL: 0.51) to the actual AUC (Arm A, SL: 0:56). Similar values suggest no inadvertent inflation.

### BC-SELECT predicted immunotherapy response better than SELECT and ENLIGHT

BC-SELECT achieved higher ROC-AUCs than SELECT and/or ENLIGHT across two immunotherapy trials (Figure 4). We were able to compare to ENLIGHT’s performance with durvalumab in breast cancer, not for pembrolizumab, as pre-computed data by ENLIGHT was available.

**Figure 4:**
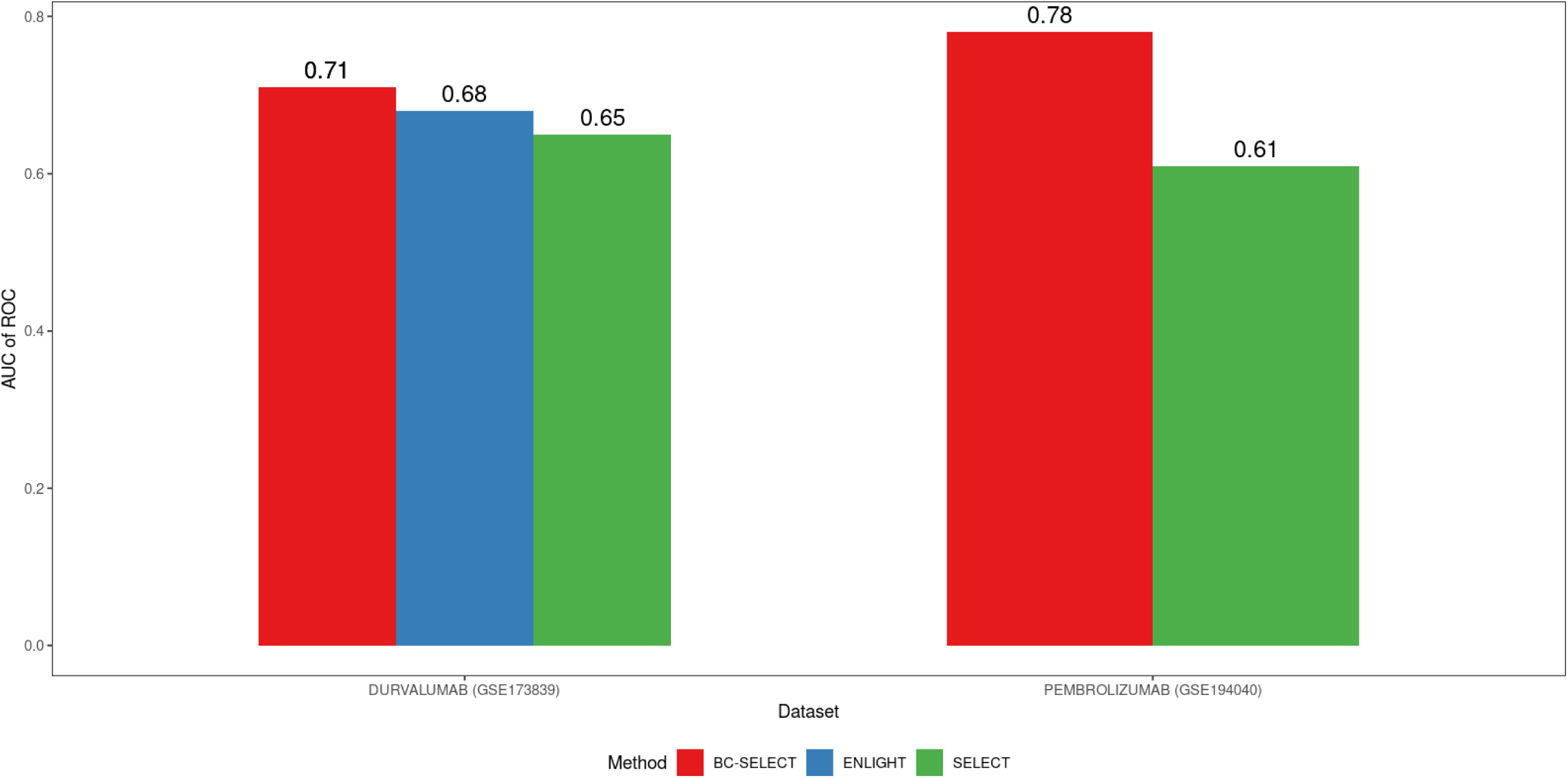
Performance comparisons between BC-SELECT and other algorithms in the context of immunotherapy. Red = BC-SELECT, Blue = ENLIGHT, and Green = SELECT. X-axis shows the specific clinical trial, and Y-axis shows the ROC-AUC. SELECT could be compared to BC-SELECT with both trials. ENLIGHT could be compared via durvalumab as this was a trial used by the developers for their validation. BC-SELECT outperforms SELECT/ENLIGHT in ROC-AUC values for both validation immunotherapy trials.

### BC-SELECT shows comparable specificity across subtypes

We assessed the specificity of the algorithm across breast cancer subtypes as possible. Among 550 patients with individual-level subtype data (347 with TNBC, 203 with HER2+), specificity was comparable, with TNBC 0.66 and HER2+ 0.64. Three trials (including both immunotherapy trials) recruited patients with HR+ disease. Specificity of BC-SELECT in HR+ disease (172 patients) was 0.76.

### Relationship between immunotherapy and PARP inhibitor BC-SELECT scores in putative BRCA-deficient tumors

To explore whether BC-SELECT scores could help distinguish between immunotherapy and PARP inhibitor benefit in tumors with putative BRCA deficiency, we compared immunotherapy-related SR scores with Veliparib-related SL/SDL scores among samples with low BRCA1/2 expression (bottom 15%) across three independent breast cancer cohorts (Figure 5). In the GSE194040^20^ Pembrolizumab cohort, higher Pembrolizumab SR scores were associated with lower Veliparib-related SL scores. A similar inverse relationship was observed between SR scores and Veliparib-related SDL scores in the GSE194040^20^ Veliparib cohort. In contrast, little association was observed in the GSE164458^33^ Veliparib cohort.

**Figure 5:**
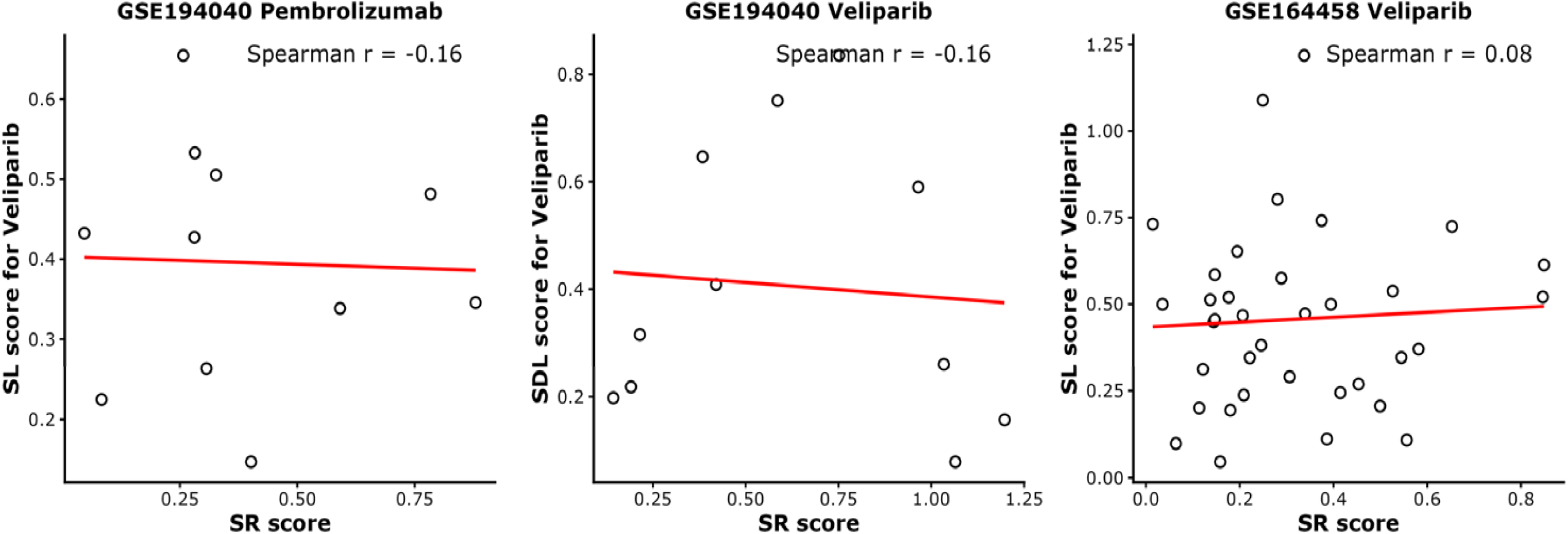
Association between immunotherapy SR scores and Veliparib-related SL/SDL scores across 3 breast cancer cohorts. To investigate potential overlap between patients predicted to benefit from immunotherapy versus PARP inhibitor, we calculated Veliparib-related SL scores in an immunotherapy-treated cohort (GSE194040 Pembrolizumab) and immunotherapy-related SR scores in PARP inhibitor-treated cohorts (GSE194040 Veliparib and GSE164458 Veliparib). Red lines represent fitted linear regression trends. Spearman correlation coefficients are displayed in each panel. SL: Synthetic Lethality. SDL: Synthetic Dosage Lethality. SR: Synthetic Rescue.

Because higher SR scores and higher SL/SDL scores are each associated with increased probability of pCR to their respective therapies, the observed inverse relationships suggest that patients predicted to benefit from immunotherapy may not necessarily be those predicted to derive the greatest benefit from PARP inhibition. Although exploratory, these findings raise the possibility that BC-SELECT scores could help prioritize treatment strategies within BRCA-deficient tumors, a setting in which clear biomarkers for selecting between immunotherapy and PARP inhibitors remain limited.

## DISCUSSION

BC-SELECT is a computational algorithm that predicts complete responders to PARP inhibitors and immunotherapy in patients with non-metastatic breast cancer. This algorithm adds to predecessors using synthetic lethality and gene expression by predicting PARP inhibitor therapy response and deepening applicability in breast cancer for multiple (potentially competing) treatment options.

BC-SELECT was developed with the goal of eventual clinical applicability. We leveraged publicly available breast cancer-specific clinical trials and datasets that included individual-level responses to targeted therapy and immunotherapy. We added patient covariates and Nanostring gene lists (to targeted therapy) to improve filtering of relevant gene partners. We used both microarray and RNA-seq data in the “training / parameter tuning” and validation steps to enable broad dataset use, though there were insufficient numbers to assess assay-specific performance. We added the “training / parameter tuning” step to improve BC-SELECT’s performance without severely limiting generalizability. Run time of BC-SELECT was within hours for large clinical trials, allowing for easy scalability as a correlative in breast cancer programs/trials. Its modular design facilitates future adaptation, where breast cancer-specific datasets with survival data can be added to the “gene-pair identification” step, and clinical trials with individual-level response data can be added to the “training / parameter tuning” step.

The intended application of BC-SELECT is prioritizing targeted therapy / immunotherapy where multiple options are available, as with non-metastatic TNBC. We sought to identify patients who may not be as responsive to approved therapies and, if eligible for multiple options, may benefit from sequential ordering of treatments in order of most likely to treat their cancer. BC-SELECT would be applied to gene expression data from tissue at the time of diagnosis and could adjust treatment with immunotherapy and PARP inhibitors as follows:

- More likely to achieve pCR with immunotherapy: Completing immunotherapy would be emphasized over other adjuvant medications
- Less likely to achieve pCR with a PARP inhibitor: PARP therapy would be deferred until after immunotherapy
- More likely to achieve pCR with a PARP inhibitor: Potential population for future trials of neoadjuvant PARP inhibitors

Germline *BRCA1/2* breast cancer is the natural application for BC-SELECT. We are actively developing collaborations for prospective data to test BC-SELECT in this population.

Another potential application in future would be hypothesis generation for potential treatments outside of current clinical guidelines. We observed stronger performance in BC-SELECT in HR+ disease relative to TNBC or HER2+. This was in the context of only three validation trials and may be an incidental observation but notably occurred with immunotherapy, where HR+ disease is notoriously resistant.

Our pathway analysis of partner genes reflected known treatment principles. Trastuzumab is given most frequently with docetaxel, a microtubule inhibitor.^23^ Pathways for veliparib included helicase and ATP hydrolyase, both involved with DNA repair.^25,34^ MHC-I downregulation increases resistance to immunotherapy.^35^ The gene families most represented among partner genes for PARP inhibitors were solute carriers (*SLC16A6, SLC27A4, SLC4A2, SLC7A14,* and *SLC7A8*) and cell cycle inhibitors. *SLC7A11* has been implicated in ferroptosis and joint mechanism of response with PARP inhibitors where *BRCA1/2* remain proficient.^36^ Five of the 10 partner genes (*CEP55, TTK, CCND3, LAG3,* and *TMEM173* (STING)) prioritized for PD-1/PD-L1 therapy have already demonstrated efficacy in studies using synthetic lethality approaches^37–43,41,42,43^. Among overlapping partner genes, co-targeting *FLT3, PLOD1, TRIB2,* or *TK1* have all shown potential increased effectiveness of PARP inhibitors or trastuzumab or both^44–48^.

BC-SELECT’s overall performance, with predictive AUCs of 0.6-0.8 in unseen trials, was consistent with -omics tests already used clinically in breast cancer. Polygenic risk scores achieve AUCs in the 0.6 range, and breast cancer gene expression assays AUCs near 0.75^49,50^. In the context we envision – prioritization of treatment options rather than eligibility where no clinical data exists - the AUCs of the algorithm fall within reasonable range for potential use.

Given our focus on genetic interactions, other predictive approaches are complementary, not competitive, with BC-SELECT. Other gene expression approaches, such as immune gene signatures or TNBC subtyping (a separate translational approach to clarify TNBCs)^51^ do not use genetic interactions. Similarly, predictors of immunotherapy response like tumor-infiltrating lymphocytes (TILs) rely on pathology.^52,53^

BC-SELECT has limitations consistent with SELECT and ENLIGHT. It requires specific drug target(s) and can only be applied to targeted therapy and immunotherapy. Many chemotherapies have numerous target effects, limiting our ability to predict chemotherapy response. BC-SELECT cannot distinguish between therapies with the same targets. BC-SELECT may not be effective if the mechanism of action does not involve synthetic lethality. This may be why we did not observe strong performance with trastuzumab in this study.

BC-SELECT was limited by data availability. We sought datasets in breast cancer with individual-level gene expression, treatment type, response, and clinical outcomes. We observed that studies with less than 25 patients total showed greater variability and reduced discriminatory power in distinguishing pCR from non-pCR. BC-SELECT was positioned to adapt as this data landscape evolves. For TNBC and HER2+, pCR as outcome was overrepresented, whereas longer-term outcomes comparable to those of HR+ gene expression tools, such as recurrence-free / progression-free / disease-free survival, were not accessible or available. We could not access publicly available individual-level treatment response data on CDK4/6 inhibitors. PIK3CA inhibitors and ESR1 inhibitors are not standard-of-care in the non-metastatic breast cancer setting. Metastatic breast cancer is the other next priority for BC-SELECT.

## CONCLUSIONS

We present BC-SELECT, a computational algorithm to predict extent of response to PARP inhibitors and immunotherapy in patients with non-metastatic breast cancer. Given the increased success rates of oncology trials studying synthetic lethality outside of PARP inhibitors,^54^ applications that use genetic interactions such as synthetic lethality to select treatments merit additional consideration. BC-SELECT demonstrates predictive performance for multiple approved therapeutic options, enabling the identification of potential complete responders and eventual decision support for sequential therapy order.

## IMPLEMENTATION

All datasets used in this paper are publicly available. Clinical trials and data collection were reported to be in accordance with the Declaration of Helsinki and National Institutes of Health Institutional Review Board (IRB) guidelines. Aggregate de-identified data was used, which was determined not to require individual patient informed consent per the National Institutes of Health IRB.

### Gene-pair identification

In BC-SELECT, we first identify pairs of genes that interact via SL, SDL, or SR, where one of the genes was a drug target. For targeted therapies, we test for SL/SDL gene pairs. DrugBank is the reference for drug target genes^55^. For immunotherapy, we test for SR pairs. *PDCD1* and *CD274* were our drug targets. We focus on targeting PD-1/PD-L1 given the availability of appropriate trial data. Pathway analysis of partner genes was performed using the Gene Ontology (Molecular Functions) database with a multiplicity-adjusted p-value of 0.05.

Gene pairs must pass the following three filtering steps in BC-SELECT: (1) A large-scale gene pair screen using *in vitro* cell-line data or *in vivo* data to select partner genes B for a given target gene A, (2) A Cox proportional hazards model in breast cancer datasets with clinical variables and overall survival to filter gene pairs with possible clinical relevance in therapy, and (3) an analysis of evolutionarily preserved relative gene expression relationships as another evidence filter. Details are provided in Figure 1, Supplementary Methods.

### Training / Parameter tuning

We use three breast cancer clinical trials with individual-level response data for training / parameter tuning: GSE66399^56^, GSE160568^57^, and NCT02489448^58^ (Supplementary Table 1). This approach avoids overfitting issues while achieving flexibility across trials. We performed a grid search to optimize the following parameters: (1) the number of significant gene pairs required to predict treatment response (ranging from 10 to 100), and (2) the odds ratio threshold to distinguish pCR versus non-pCR.

For *targeted therapy*, we select the top 55 SL/SDL gene pairs, with ranking based on adjusted significance from Cox proportional hazards. We used only either SL or SDL depending on predictive performance to avoid mixed direction of effect. For *immunotherapy*, we selected the top 10 SR gene pairs, with ranking based on phylogenetic similarity scores. We used an odds ratio threshold of greater than 0.51 for “pCR.”

**Note**: We include the “training / parameter tuning” step to optimize the parameters of significant gene pairs and odds ratio threshold across the tested therapies. While this step improved validity, it is not strictly required. The gene pairs fed into “training / parameter tuning” from the “gene pair identification” step are already derived from large-scale datasets with intensive filtering.

### BC-SELECT predicted response score

The predicted response score in BC-SELECT was the product of the **partner gene expression ratio** and the **drug target gene expression level**. How we calculate each element is described in detail in the Supplementary Methods.

The breast cancer-specific and immune-specific gene lists used as part of the partner gene expression ratio calculation were selected through commercial Nanostring panels.^59,60^ The two Nanostring gene lists are provided in Supplementary Table 3.

### Validation trials

We used publicly available phase II to III clinical trials with at least 25 patients with stage I-III breast cancer, individual level gene expression data, and reported treatment response. We selected pCR to measure response, as this treatment response variable could be harmonized across clinical trials. We used pCR as reported by the clinical trials – a binary measure of either 0 (residual disease/non-pCR, including RCB levels) or 1 (complete response). All trials used the definition of pCR as the absence of residual invasive tumor in the breast and axillary lymph nodes^61^. Treatment response for these trials served solely as the “ground truth” comparator versus BC-SELECT’s predicted response score.

We included a total of 9 clinical trials, with 7 across 3 targeted therapies (trastuzumab, olaparib, veliparib)^20,33,62–66^ and 2 clinical trials for immunotherapy^20,66^ (Supplementary Table 1). Where trials recruited HR+ patients, evaluation of these patients was handled in the same manner as for the TNBC and HER2+ patients.

### Validation of treatment response prediction

We compared BC-SELECT predicted response scores per patient to the actual responses reported in each trial. We calculated odds ratios with 95% confidence intervals and p-values using Fisher’s exact test, as well as ROC-AUC and precision-recall curves to assess prediction performance.

**Note:** GSE173839^66^ involved treatment with two drugs in our study (olaparib and durvalumab). We independently evaluated the predicted response of olaparib and durvalumab and generated two separate BC-SELECT scores, one for each drug, and two separate AUCs.

### Cross-therapy score association analysis

To determine whether BC-SELECT identifies similar or distinct patient populations for immunotherapy and PARP inhibition, we examined the relationship between immunotherapy-related SR scores and Veliparib-related SL/SDL scores. Because germline *BRCA1/2* mutation status was unavailable for our datasets, we used *BRCA1* and *BRCA2* expression as a proxy for BRCA deficiency. Samples were ranked by the average expression of *BRCA1* and *BRCA2* in each cohort, and the bottom 15% of samples were selected for analysis. For the immunotherapy-treated GSE194040^20^ Pembrolizumab cohort, Veliparib-related SL scores were calculated using pre-treatment gene expression data using the previously identified Veliparib SL partner genes. Conversely, for the PARP inhibitor-treated GSE194040^20^ Veliparib and GSE164458^33^ Veliparib cohorts, immunotherapy-related SR scores were calculated using the top 10 SR partner genes identified for immunotherapy analysis. Associations between the two scores were evaluated using Spearman rank correlation, given the modest sample sizes and the possibility of non-linear relationships between the scores.

### Statistical analysis

For *targeted therapy*, we perform a Wilcoxon rank-sum test to rank and select gene pairs. For *immunotherapy*, we perform a hypergeometric test to identify pairs of genes observed as under-expressed more than expected by chance in breast cancer datasets. We use a Cox proportional hazards model as part of the gene pair filtering. More details on this testing are described in the Supplementary Methods.

We estimated the statistical significance of BC-SELECT’s predicted response score and components compared to chance using Monte-Carlo simulations of the ROC-AUC (Supplementary Figure 2, Supplementary Methods).

All analyses were conducted using R v.4.2. To control the false discovery rate, we apply the Benjamini–Hochberg procedure (BH step-up procedure) as a multiplicity correction.

## Project Name

BC-SELECT

## Project Home page

https://github.com/yewon-kim76/BC-SELECT

## Operating system

Platform independent

## Programming language

R

## Other requirements

R libraries called in software

## License / Restrictions

This software is created by U.S. federal government employees and is part of the public domain.

## List of Abbreviations

## Ethical Approval and Consent to Participate

Clinical trials and data collection for the public datasets used in this paper were reported to be in accordance with the Declaration of Helsinki and National Institutes of Health Institutional Review Board (IRB) guidelines. Aggregate de-identified data was used, which was determined not to require individual patient informed consent per the National Institutes of Health IRB.

## Consent for Publication

N/A

## Data Availability Statement

The data analyzed during the current study are available in the publicly available repositories as described in the Supplementary Files. The underlying code for BC-SELECT is available on GitHub.

## Author contributions

**Yewon Kim**: Methodology, Software, Validation, Analysis, Writing - Original And Revision. **Matthew Nagy**: Software, Validation, Writing – Revision. **Becca Pollard**: Data Curation. **Padma Sheila Rajagopal**: Conceptualization, Methodology, Validation, Writing - Revision, Resources, Data Curation, Supervision, Administration, Funding.

### Acknowledgments

We wish to thank Dr. Eytan Ruppin, Dr. Lipika Ray, and Dr. Nishanth Nair for their support and counsel throughout the drafting of this manuscript. We also wish to thank Dr. Kim Blenman, Dr. Lajos Pusztai and Dr. Daniel Stover for their generosity in providing individual-level training data or data-use recommendations in support of this project. This research was supported by the Intramural Research Program of the National Institutes of Health (NIH). The contributions of the NIH author(s) were made as part of their official duties as NIH federal employees, are in compliance with agency policy requirements, and are considered Works of the United States Government. However, the findings and conclusions presented in this paper are those of the author(s) and do not necessarily reflect the views of the NIH or the U.S. Department of Health and Human Services. This work utilized the computational resources of the NIH HPC Biowulf cluster. (http://hpc.nih.gov)

